# Bisphenol A affects the development and the onset of photosymbiosis in the acoel *Symsagittifera roscoffensis*

**DOI:** 10.1101/2024.05.03.592490

**Authors:** R. Pennati, N. Cartelli, C. Castelletti, F. Ficetola, X. Bailly, S. Mercurio

**Affiliations:** Department of Environmental Science and Policy, Università degli Studi di Milano, Italy; Multicellular Marine Models (M3) team, Station Biologique de Roscoff, CNRS/Sorbonne Université. Place Georges Teissier, 29680 Roscoff. France

**Keywords:** photosymbiosis, photosymbiogenesis, BPA, miR-124, algae, worm, embryogenesis

## Abstract

Photosymbiosis indicates a long-term association between animals and photosynthetic organisms. It has been mainly investigated in photosymbiotic cnidarians, while other photosymbiotic associations have been largely neglected. The acoel *Symsagittifera roscoffensis* lives in obligatory symbiosis with the microalgal *Tetraselmis convolutae* and has recently emerged as alternative model to study photosymbiosis. Here, we investigated the effects of Bisphenol A, a common plastic additive, on two pivotal stages of its lifecycle: aposymbiotic juvenile development and photosymbiogenesis. Based on our results, this pollutant altered the development of the worms and their capacity to engulf algae from the environment at concentrations higher than the levels detected in seawater, yet aligning with those documented in sediments of populated areas. Data provide novel information about the effects of pollutants on photosymbiotic associations and prompt the necessity to monitor their concentrations in marine environmental matrices.

## 1. Introduction

Photosymbiosis is a widespread phenomenon observed in multiple animal phyla, including sponges, cnidarians, acoelomorphs, platyhelminths, mollusks, ascidians, and even vertebrates (Melo Clavijo et al., 2018; Venn et al., 2008). The term is used to indicate a long-term association between a heterotrophic organism and a phototrophic one (Cowen, 1988; Yonge, 1934). The symbioses between animals and algae are the results of a precise balance involving different physiological and environmental factors, and are thus particularly susceptible to anthropogenic stressors. The peculiar nature of this relationship has boosted many scientific investigations, most of which have focused on the effects of environmental perturbations and pollutants on corals and their *Symbiodinium* symbionts (Bielmyer et al., 2010; Kuzminov et al., 2013; Marzonie et al., 2021; van Dam et al., 2015; Venn et al., 2008). It has been reported that some compounds can trigger bleaching events, i.e. the loss of algae from the symbiosis. Cnidarians can lose their symbionts when exposed to several pollutants, such as heavy metals and herbicides (Bielmyer et al., 2010; Kuzminov et al., 2013; Marzonie et al., 2021), and additive effects from chemicals and environmental stressors have been also demonstrated, (van Dam et al., 2015). Similar studies are instead rare for other photosymbiotic associations, which however play a key role in many marine ecosystems (Erwin and Thacker, 2007; Not et al., 2016; Venn et al., 2008). Furthermore, most research has studied the effects of stressors on bleaching processes, while their impact on the development of aposymbiotic embryos as well as on the onset of photosymbiosis have been largely neglected probably due to difficulties in the experimental design with common cnidarian models. Indeed, controlling life cycle in captivity offers a valuable access to any developmental stages along the species lifespan. For photosymbiosis, this additionally implies to cultivate the photosynthetic partner for inducing the photosymbiosis when the juvenile is aposymbiotic i.e. when there is no vertical transfer of the symbiont.

In the last decade, the photosymbiotic acoel (phylum xenacoelomorpha) *Symsagittifera roscoffensis* and its algal symbionts *Tetraselmis convolutae* have emerged as alternative biological system in photosymbiosis research (Arboleda et al., 2018; Bailly et al., 2014). *S. roscoffensis* is a marine acoel belonging to the Saggitiferidae family that lives in obligatory symbiosis with the green microalgal chlorophyte *T. convolutae. S. roscoffensis* is a gregarious species forming large populations on sandy shores of the North Atlantic coasts (Bailly et al., 2014). During the breeding season, hermaphroditic gravid adults deposit embryos inside mucilaginous protective capsules called cocoons. Embryogenesis lasts about 4-5 days and results into colourless aposymbiotic juveniles. The symbiont transmission is strictly horizontal; hatching juveniles ingest environmental free microalgae within the first days and keep them for their lifetime. Algae are internalized into vacuoles in the digestive syncytium where they lose their cell wall, flagella and eyespot, thus deeply changing their phenotype. Microalgae proliferate so that adult worms finally host around 120,000 algae, forming a dense layer underneath the body wall and giving a green colour to the animals. Adult worms do not eat any food and depend entirely on organic supply from microalgal photosynthetic products (no mixtrophy regime has been to date demonstrated). Despite very few data on the trophic relationship, *T. convolutae* uses the uric acid, a nitrogen waste product produced by the worm, as an endogenous nitrogen source (Bailly et al., 2014; Boyle and Smith, 1975; Doonan and Gooday, 1982; Douglas, 1983).

The handiness of these organisms, together with the optimized laboratory conditions and the recently provided genomic resources have made this species an excellent model to design functional exploration experiments and investigating many biological questions (Bailly et al., 2014; Martinez et al., 2022). In this work, we used the *S. roscoffensis*/*T. convolutae* association to study the effects of the common plastic additive, Bisphenol A (BPA), on embryogenesis and photosymbiogenesis.

BPA is one of the chemicals with the highest annual production; in 2021 it reached 11 million tons worldwide and its global demand is expected to increase significantly in the future (Devi et al., 2023). Thus, BPA is continuously released in the environment and, despite its short half-life, it is considered as a pseudo-persistent chemical (Flint et al., 2012). BPA is, indeed, one of the most common plastic additives recovered in the marine environment. It is generally mixed with plastic polymers during the manufacturing processes (Hermabessiere et al., 2017), but it can easily migrate from plastic products to the medium in contact with it (Brede et al., 2003; Geens et al., 2010; Hahladakis et al., 2018). This chemical reaches the marine environment through different routes, including wastewater, river transport or leachate from plastic items, and they accumulate differently in marine compartments (Hermabessiere et al., 2017; Li et al., 2024). In seawaters, BPA has been generally detected in traces (ng/l) but it could reach higher concentrations (µg/l) in polluted areas, such as Singapore coastal water (Basheer et al., 2004; Corrales et al., 2015; Hermabessiere et al., 2017). The occurrence of this chemical in sediments and biota is source of concern as well: due to its hydrophobic properties, BPA may accumulate in these environmental matrices, reaching warning concentrations at local scale (Flint et al., 2012; Gao et al., 2023; Hermabessiere et al., 2017; Xu et al., 2015). Adverse effects of BPA have been reported in different marine species and include impairment of animal development and reproductive processes. For examples, BPA exposure affected sexual dimorphism in the marine medaka *Oryzias melastigma* (Yamamoto et al., 2023) and induced expression of the vitellogenin-like protein and spawning event in the mussel *Mytilus edulis* (Aarab et al., 2006). During invertebrate development, effects include cleavage arrest, embryonic malformations, such as neural defects, and metamorphosis inhibition (Mansueto et al., 2011; Mercurio et al., 2022; Messinetti et al., 2019, 2018; Miglioli et al., 2021; Zhou et al., 2011).

Here, we took advantages from the unconventional photosymbiotic model *S. roscoffensis* to investigate the still unexplored impact of this widespread contaminant on two important biological processes: the development of marine aposymbiotic embryos and the onset of photosymbiosis in order to evaluate BPA possible interference with these complex processes.

## 2. Materials and methods

### 2.1. Animals

Adults of *S. roscoffensis* were collected at low tide from shores near Roscoff (NW France) in late October. They were immediately transferred in incubators and maintained in filtered seawater (FSW) at 14 ± 1°C with a photoperiod set at 10 h:14 h (light:dark). After about 48 hours, gravid specimens naturally spawn: cocoons were immediately collected and kept in FSW in dark conditions, to be used in the following experiments. *T. convolutae* culture was obtained from the Roscoff Culture Collection.

### 2.2. Embryo exposures

*S. roscoffensis* embryos within their cocoons at early cleavage stages (2-cell and 4-cell stage) were collected under a stereomicroscope and exposed to increasing concentrations of BPA (0.05, 0.1, 0.5, 1, 5, 10, µM). BPA (MW = 228.29) was purchased from Sigma (Milan, Italy). A stock solution of 100 mM BPA was prepared dissolving 22.8 mg of powder in 1 ml of dimethyl sulfoxide (DMSO). Tested concentrations were obtained by scale dilutions in FSW and half of the exposure volume was renewed with freshly prepared solutions every day. BPA exposure levels were defined based on published works (Hermabessiere et al., 2017; Mercurio et al., 2022; Messinetti et al., 2019, 2018; Miglioli et al., 2021) and preliminary trials. Two control groups were set up: CO embryos were reared in FSW and DMSO animals were maintained in FSW with 0.01% DMSO, in order to verify possible solvent effects. All exposures were performed in triplicates at 14 ± 1°C in glass Petri dishes and lasted 5 days, i.e. when aposymbiotic juveniles hatched from their cocoons. At the end of the experiments, samples were observed under a stereomicroscope to score the number of free-swimming individuals, unhatched individuals still inside their cocoons (showing a delay of development) and dead embryos. Then, embryos were relaxed in a solution of 7% MgCl_2_ in FSW for 10 minutes and fixed for 1 h at room temperature in a solution of 4% paraformaldehyde, 0.5 M NaCl and 0.1 M 3-(N-morpholino)propanesulfonic acid (MOPS fixative; pH 7.5). After several rinsing in Phosphate Buffered Saline (PBS) with 0.1% Tween-20 (PBT), samples were photographed under a Leica optical microscope and then dehydrated and stored in 70% ethanol at - 20°C.

### 2.3. In situ hybridization

The effects of BPA on the development of the nervous system were analyzed performing *in situ* hybridization experiments with a commercial DIG-labelled Locked Nucleic Acid probe (LNA; Exiqon, Norway) against the neuro-specific microRNA miR-124. The sequence of the LNA probe (5’-TTGGCATTCACCGCGTGCCTTA -3’) was designed complementary to the *S. roscoffensis* miR-124 sequence previously reported (Sempere et al., 2007). A protocol of whole mount *in situ* hybridization with LNA probes was specifically optimized modifying those used for other marine invertebrates (Mercurio et al., 2020, 2019a, 2019b). Briefly, samples fixed in MOPS fixative were rinsed several times in PBT and hybridized with LNA probe at 55°C in 50% formamide, 5x SSC, 100 μg/ml yeast RNA; 50 μg/ml Heparin; 0.1% Tween-20 for 5 days. To eliminate the probe excess, samples were washed several times in a solution of 50% formamide, 5x SSC, 0.1 Tween-20 at 55°C and then rinsed in PBT at room temperature. Samples were pre-incubated for 2 h in blocking solution (25% deactivated goat serum in PBT) and incubated overnight at 4°C in blocking solution with anti-DIG antibody conjugated to alkaline phosphatase (Roche Diagnostics, Germany; 1:2000). After several washes in PBT, staining reaction was carried out in alkaline phosphataselabeled buffer (100 mM NaCl, 100 mM Tris HCl, pH 9.5, 50 mM MgCl_2_, 0.1% Tween□20) + 2.3 μl/ml 4□Nitrotetrazolium Blue chloride and 3.5 μl/ml 5□Bromo□4□chloro□3□indolyl phosphate p□toluidine salt in dark conditions at room temperature. When a satisfactory signal was developed, the reaction was stopped by rinsing the samples in PBT and fixing them in MOPS fixative for 1 h. Finally, hybridized juveniles were mounted in 80% glycerol and photographed under an optical microscope equipped with Leica DFC□320 Camera.□

### 2.4. Co-exposures of algae and juveniles

To investigate BPA impact on photosymbiogenesis, we co-exposed *T. convolutae* microalgae and aposymbiotic juveniles of *S. roscoffensis* to BPA tested concentrations (controls, 0.05, 0.1, 0.5, 1, 5, 10, µM BPA). To mimic an environmental scenario, algae and juveniles were separately exposed to BPA for 12 hours prior to mix them in the same exposure solution. Algal pre-treatment involved mixing 5 ml of a dense *T. convolutae* culture with an equal volume of a solution containing BPA concentration twice that of the exposure level. Similarly, 3 ml of FSW containing aposymbiotic juveniles were mixed with an equal volume of 2x concentrated solution of BPA to obtain the final tested solution. Then, co-exposures were carried out added BPA pre-treated juveniles to their corresponding BPA-treated algae culture. Experiments lasted 72 h and were performed in triplicates. *S. roscoffensis* juveniles were then washed in FSW, fixed in MOPS fixative for 1 h at room temperature and stored at 4°C.

### 2.5. Evaluation of symbiogenesis

The effects of algae and juvenile co-exposure to BPA were determined by counting the number of algae that were recruited by each treated juvenile. Exposed juveniles were mounted in 80% glycerol and observed under a fluorescence microscope equipped with red filter, i.e. transmitting light with a wavelength between 550–700 nm. Taking advantage of the endogenous autofluorescence produced by the chlorophyll (Fricker and White, 1992; Yentsch and Menzel, 1963; Zhang et al., 2010) we could easily count algae internalized by each *S. roscoffensis* juvenile and annotate algae number/sample for each experimental group.

### 2.6. Statistical analysis

All the analyses were performed in the R 3.6.3 environment (R Core Team, 2019). Analysis of variance (ANOVA), followed by honestly significant difference Tukey’s post hoc test, was performed to assess the effect of BPA on embryonic development as previously reported (Mercurio et al., 2021). We used generalized mixed models (GLMM) to assess the relationship between BPA concentration and the number of symbionts, while taking into account potential differences between experimental trials. Models were run in the glmmTMB package using a negative binomial error distribution to take into account overdispersion of counts across individuals within treatment (Brooks et al., 2017). Tukey’s post-hocs were then run using the multcomp package to identify differences between treatments (Hothorn et al., 2008). Binomial generalized linear models with probit link function (GLM) was used to assess the impact of different concentrations of BPA on 1) mortality rate of embryos and 2) the proportion of embryos with malformations within alive embryos,. Due to overdispersion, for GLMs we used a quasibinomial error and calculated significance using a F test (Venables and Ripley, 2002). On the basis of GLMs, we calculated LC_50_ (median lethal concentration) – EC_50_ (median effective concentration) i.e. the concentration of PBA that determines 50% mortality / 50% of embryos with developmental alterations.

## 3. Results

### 3.1. Embryogenesis

BPA effects on the embryogenesis of *S. roscoffensis* were evaluated by comparing the morphology and the percentages of swimming juveniles, unhatched embryos, which were still inside their cocoons after the exposure period, and dead samples among the experimental groups.

Most of controls and solvent control juveniles (DMSO) regularly hatched 5 days post cocoon laying (CO: 94.7%; DMSO: 97.9%; Table 1). They displayed an elongated shape with a broad anterior extremity and a narrow posterior one (Fig. 1 A and B). The anterior sensory organ, the statocyst, was well-developed and samples moved actively crawling on the bottom of the petri dishes. From 1 μM BPA, the incidence of unhatched embryos significantly increased compared to controls: 37.5% of the exposed embryos remained inside their cocoons at the end of the exposure period (Tukey HSD Post-hoc Test; BPA *vs* CO and vs DMSO: *P*<0.001; Fig. 1 F; Table 1). Although these embryos within their cocoons exhibited movements, they showed developmental delayed or were incapable of hatching. Compared to the control elongated phenotype, most of them showed a rounded morphology as body elongation had not occurred yet. Hatched juveniles were similar to controls and normally crawled on the substrate. At 5 μM BPA, more than 95% of embryos were still inside their cocoons (Tukey HSD Post-hoc Test; 5 μM BPA *vs* CO/DMSO: *P*<0.001; Fig. 1 G; Table 1). The embryos were roundish and the two body extremities were not distinguishable (Fig. 1 F). Statocysts were not always recognizable. The highest tested concentration, 10 μM BPA, resulted lethal for the totality of the exposed samples (Tukey HSD Post-hoc Test; 10 μM BPA *vs* CO/DMSO: *P*<0.001; Fig. 1 G, Table 1) and developmental disruption occurred early during embryogenesis as dead embryos were already observable after 24 h of exposure. Probit analysis (Fig. 1 H) confirmed these results: LC_50_ was 6.31 μM (95% CI for the coefficient estimate: 0.318-3.950; *F*_*1,19*_ = 7.18, *P* = 0.015) and EC_50_ was 1.28 μM (95% CI for the coefficient estimate: 0.382-3.192; *F*_1,16_= 8.27, *P* = 0.011). BPA teratogenic index (TI = LC_50_/EC_50_) was 4.93.

**Table 1.** Percentages of swimming juveniles, unhatched and dead embryos among the experimental groups. Values are expressed as means ± standard errors.

|  | **Swimming juveniles (%)** | **Unhatched embryos (%)** | **Dead embryos (%)** |
| --- | --- | --- | --- |
| CO | 94.7 ± 2.5 | 5.3 ± 2.5 | 0 ± 0 |
| DMSO | 97.9 ± 1 | 2 ± 1 | 0 ± 0 |
| 0.1 µM BPA | 93.7 ± 5 | 1.6 ± 0.8 | 4.6 ± 4.6 |
| 0.5 µM BPA | 98.2 ± 0.9 | 1.8 ± 0.9 | 0 ± 0 |
| 1 µM BPA | 57 ± 11 | 37.5 ± 5.6 | 5.4 ± 5.4 |
| 5 µM BPA | 0 ± 0 | 95.24 ± 4.7 | 4.7 ± 4.7 |
| 10 µM BPA | 0 ± 0 | 0 ± 0 | 100 ± 0 |

**Figure 1.**
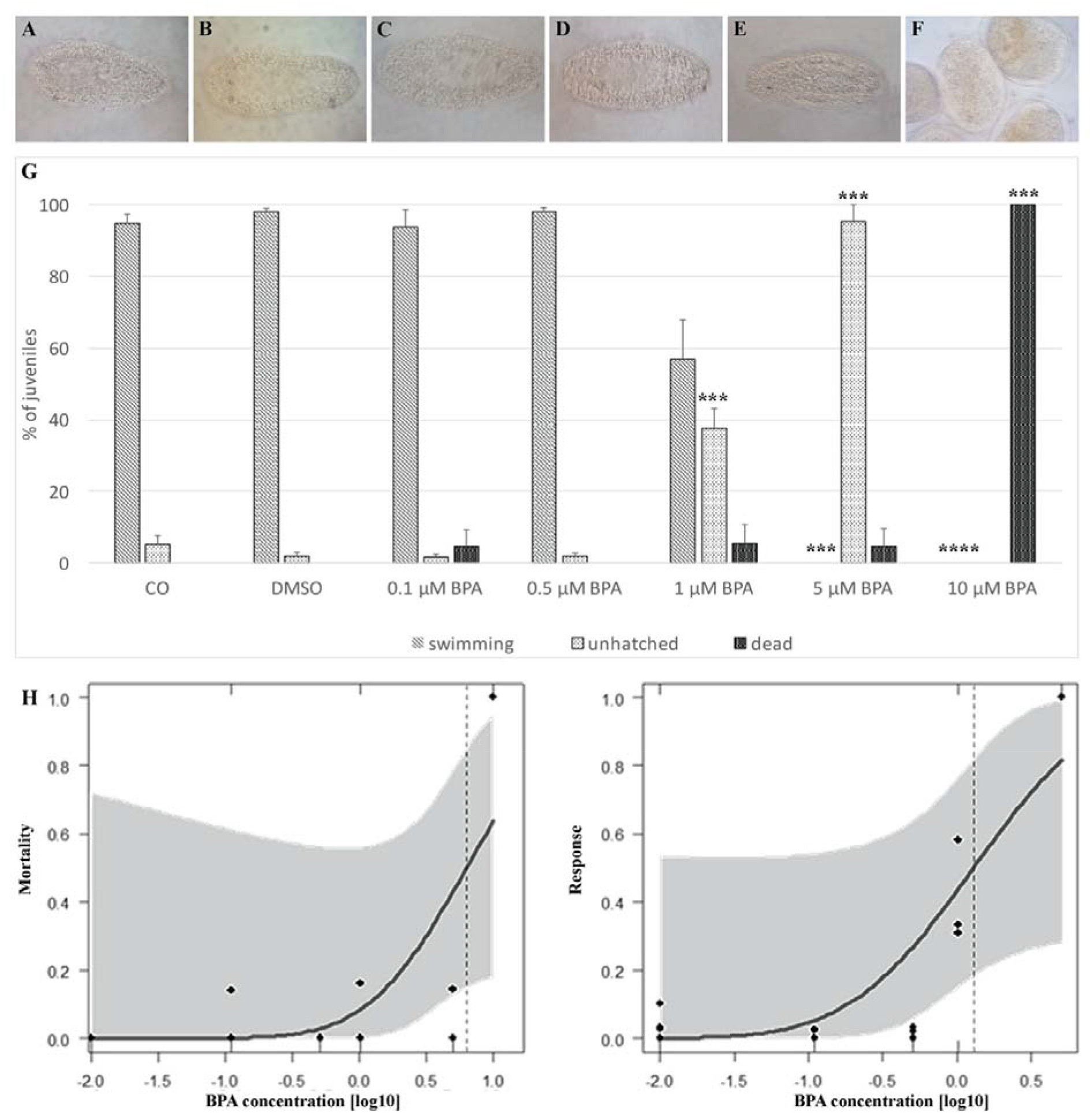
Effect of BPA exposure on the development of *S. roscoffensis*. **A-F**) Morphology of controls, CO (**A**) and DMSO (**B**) and of juveniles exposed to 0.1 µM (**C**), 0.5 µM (**D**), 1 µM (**E**) and 5 µM (**F**) BPA. Scale bar = 50 µm. **G**) Percentages of swimming, unhatched and dead juveniles exposed to BPA expressed as mean values and standard errors; differences from controls: the repetition of each symbol indicates the level of significance: *** p < 0.001. H) BPA dose-response curves for mortality and developmental alterations determined by probit analysis. Continuous blue line indicates the mortality/alteration rates predicted by models; black dots indicate the observed mortality/alteration rates in the different experimental replicates, shaded areas are 95% CIs. The vertical dotted lines indicate the LC_50_ and EC_50_ values.

### 3.2. Neural development

The effects of BPA exposure on the development of nervous system were investigated by analyzing the expression of miR-124, a pan-neuronal microRNA (Fig. 2). In control juveniles, miR-124 expression was detected in the central nervous system, in the neuropile surrounding the statocyst (Fig. 2 A). Similar expression was observed in juveniles exposed to low concentrations of BPA (up to 1 μM BPA; Fig. 2 B and C). Conversely, miR-124 was not detectable in samples exposed to 5 μM BPA (Fig. 2 D).

**Figure 2.**
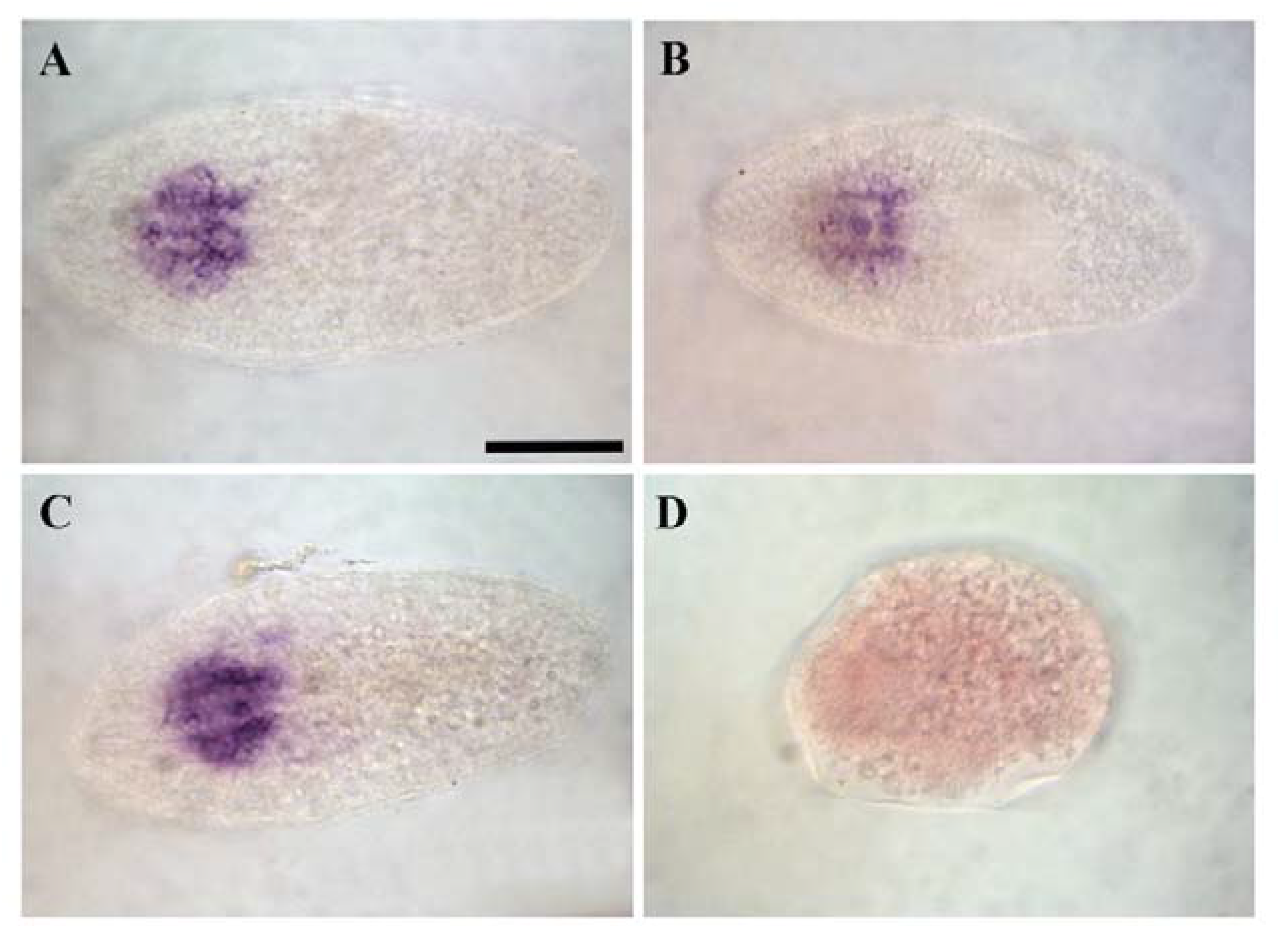
Expression of miR-124 in *S. roscoffensis* juveniles exposed to different concentrations of BPA. A) DMSO; B) 0.5 µM BPA; C) 1 µM BPA; D) 5 µM BPA. Scale bar = 50 µm

### 3.3. Photosymbiogenesis

*S. roscoffensis* aposymbiotic juveniles and *T. convolutae* algae were co-exposed to increasing concentrations of BPA to assess its effects on the onset of photosymbiosis. The number of microalgal photosymbionts showed a very strong decrease at growing BPA concentrations (GLMM with negative binomial models: 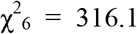, *P* < 0.0001; Fig. 3). The number of symbionts remained roughly constant (average: 9-11 symbionts per individual) at concentrations up to 1 μM (Fig. 3 A and B), with no significant changes compared to controls (Tuckey’s post-hoc: *P* > 0.5 for all pairwise comparisons; Fig. 3 E). At 5 μM (Fig. 3 C), the average number of algae dropped at 1.3, with a significant decrease compared to both the control and all the lower tested concentrations (Tuckey’s post-hoc: all *P* < 0.001; Fig. 3 E). The average number of algal symbionts further dropped at 10 μM (average: 0.5 symbionts / individual; Fig. 3 D), with a significant decrease compared to 5 μM (*P* = 0.01) and to all the previous concentrations (all *P* < 0.001; Fig. 3 E).

**Figure 3.**
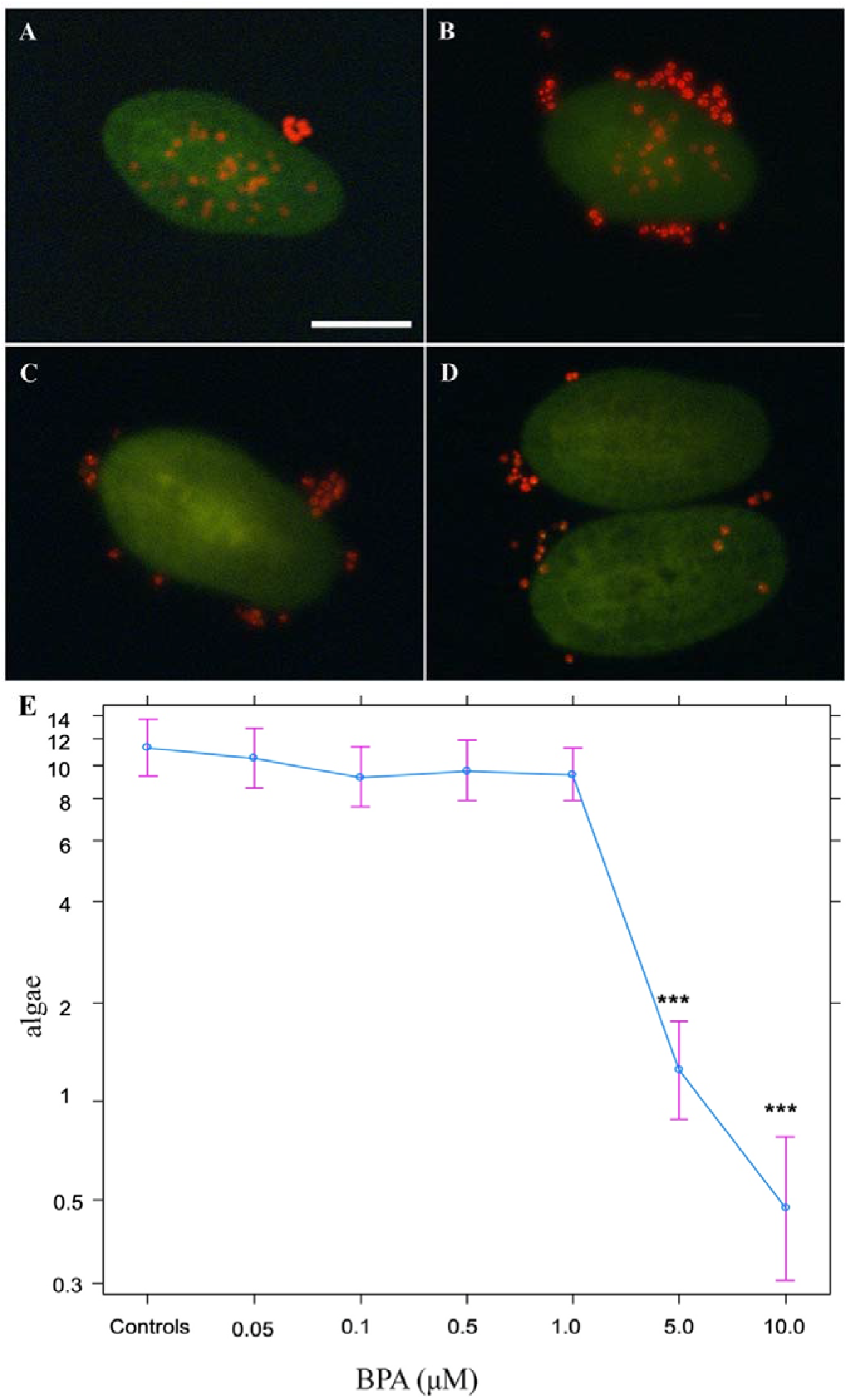
Evaluation of BPA effects on the onset of photosymbiosis. **A-C**) Juveniles of *S. roscoffensis* juveniles (green) and *T. convolutae* algae (red) observed under a fluorescence microscope. Free-living algae attached on the outer surface of the animals are often visible. Controls exposed to 0.01% DMSO (**A**) and samples exposed to 1 µM BPA (**B**) displayed numerous internalized algae in their body (∼20). In samples exposed to 5 µM (**C**) and 10 µM (**D**) algal symbionts were rare (<5). Scale bar = 50 µm. **E**) Variation in the average number of photosymbionts across BPA concentrations. Error bars are 95% confidence intervals, estimated on the basis of generalized mixed models. *** indicate significant differences (*P* < 0.001) compared to the controls.

## 4. Discussion

In the present work, we exploited the emerging photosymbiotic model organism *S. roscoffensis* to study the effects of BPA, a common marine pollutant (Corrales et al., 2015; Flint et al., 2012; Hermabessiere et al., 2017), on its embryonic development and photosymbiogenesis, revealing that this chemical can disrupt both processes.

BPA induced developmental delay and alterations starting from less than 1 μM (22.800 ng/L) (EC_50_ was 1.28 μM). Similarly, its effects on development, reproduction, settlement and metamorphosis in 3 coral reef photosymbiotic invertebrates, *Stylophora pistillata, Millepora dichotoma*, and *Rhytisma fulvum*, have been demonstrated to be limited at environmentally relevant concentrations (Vered and Shenkar, 2022). However, these concentrations are higher than the levels detected in seawater, yet aligning with those documented in sediments of populated areas (Corrales et al., 2015; Hermabessiere et al., 2017). Sediments are the environmental matrix with the highest concentration of BPA (Torres-García et al., 2022) where concentration as high as 270 ng/g dw have been reported (Dan Liu et al., 2017). *S. roscoffensis* is a benthic invertebrate which lives in close association with the sandy substrate (Bailly et al., 2014), and thus could experience high level of hydrophobic pollutants, such as BPA, which accumulate in this environmental compartment. This result is particularly concerning as this photosymbiotic acoel has an important ecological role. *S. roscoffensis* has a highly efficient photosynthetic ability and its large populations of the intertidal zone can generate a primary production close to those reported for coral reefs (Androuin et al., 2020; Thomas et al., 2023). Moreover, this species has been proposed as a possible natural bioremediator as it shows high nitrate assimilation rates, and it can thus act as an important *in situ* nitrate recycler in polluted marine environments (Carvalho et al., 2013).

At the high tested concentrations (from 5 µM), BPA affects worm morphology as most juveniles appeared roundish and statocysts were not always recognizable. Moreover, thanks to a newly optimized protocol for whole mount *in situ* hybridization, we reported for the first time the expression of the pan-neural miRNA, miR-124, in acoels and demonstrated that BPA disrupts neural development in *S. roscoffensis*. Similar effects have been already documented in other marine invertebrates. In the ascidians *Ciona robusta* and *Phallusia mammillata*, BPA, at concentrations higher than 5 µM, impaired sensory organ formation and central nervous system development (Messinetti et al., 2019, 2018). It interfered with the differentiation of GABAergic and dopaminergic neurons in *C. robusta* (Messinetti et al., 2019) while serotoninergic system was the main neural target in the mussel *Mytilus galloprovincialis* (Miglioli et al., 2021). However, EC_50_ and LC_50_ are highly variable among marine organisms: in *C. intestinalis* and *C. robusta*, BPA EC_50_ resulted 8.25 μM (1.88 mg/l) and 7.04 μM (1.6 mg/l) while BPA LC_50_ was 13.42 μM (3.06 mg/l) and 9.36 μM (2.13 mg/l) respectively (Mercurio et al., 2022); LC_50_ was 107.2 mg/l for the zooplanktonic grazer *Artemia salina*, 11.5 mg/l for the snail *Heleobia australis*, and 3.5 mg/l for the fish *Poecilia vivipara* (Naveira et al., 2021b). Comparing to these data, *S. roscoffensis* appeared more sensitive to BPA than many other animals: EC_50_ was 1.28 μM (0.29 mg/l) and LC_50_ was 6.31 μM (1.4 mg/l).

The onset of photosymbiosis was affected by BPA exposure as well: growing BPA concentrations induced a drastic decrease in algal content in aposymbiotic juveniles, suggesting that the chemical interfered with some of the mechanisms involved in the photosymbiogenesis. The bleaching phenomenon has been extensively investigated in coral and their symbionts (Bielmyer et al., 2010; Kuzminov et al., 2013; Marzonie et al., 2021; van Dam et al., 2015; Venn et al., 2008), in which loss of algae can be triggered by different pollutants, such as heavy metal and herbicides (Bielmyer et al., 2010; Kuzminov et al., 2013; Marzonie et al., 2021). On the contrary, the mechanisms involved in photosymbiogenesis are mostly unexplored. Thanks to *S. roscoffensis* / *T. convolutae* biological system and its optimized experimental conditions (Bailly et al., 2014), we explored for the first time BPA interference with the onset of photosymbiotic interaction. Data related to EC_50_ and LC_50_ indicate that *Tetraselmis sp*. algae well-tolerate a wide range of BPA concentrations (Naveira et al., 2021a), while our Probit results suggested that *S. roscoffensis* acoels are sensitive to the contaminant (EC_50_=1.28 μM; LC_50_=6.31 μM). We could thus hypothesize that BPA effects on photosymbiogenesis were mainly determined by disruptions in the interaction mechanisms exerted by the worms, but the current lack of information about cues controlling symbiotic interactions prevents any more specific considerations.

As these concentrations are higher than the average environmental ones, this result appeared quite comforting, but more awareness is necessary. In fact, concentrations of the same magnitude of EC_50_ of *S. roscoffensis* have already been reported in both surface water and groundwater (25,000 and 30,000 ng/L BPA respectively) in the Santa Catarina River, Mexico (Cruz-López et al., 2020) and in sludge from sewage treatment works (16300 ng/L BPA) in UK (Petrie et al., 2019). Moreover, marine animals are exposed simultaneously to a variety of pollutants, which may affect photosymbiosis at different levels and can cause mixture additive effects (Altenburger et al., 2003) and thus even at low concentrations might be able to disrupt this algal-animal relationship.

Photosymbiotic associations are frequent in marine environment and involved a great variety of organisms, most of which plays a key ecological role (Decelle et al., 2015; Erwin and Thacker, 2007; Not et al., 2016; Venn et al., 2008). However, so far experimental investigations have been mainly focused on photosymbiotic corals and anemones (cnidarian/dinoflagellate associations) due to their ecological importance for coral reef ecosystems (Bielmyer et al., 2010; Kuzminov et al., 2013; Marzonie et al., 2021; Venn et al., 2008). Here, we exploited the alternative model *S. roscoffensis* to study BPA effects on two focal developmental stages of marine photosymbionts. Our results provided novel information about one of the most common pollutants (Corrales et al., 2015; Flint et al., 2012; Hermabessiere et al., 2017), which can help shape effective science-driven policies for environmental management. Moreover, we provided new experimental tools to design functional experiments and investigate biological questions related to photosymbiosis and the molecular interactions involved in its onset.

